# Neuronal Odor Coding in the Larval Sensory Cone of *Anopheles coluzzii*: Complex Responses from a Simple System

**DOI:** 10.1101/2020.09.09.290544

**Authors:** Huahua Sun, Feng Liu, Adam Baker, Laurence J. Zwiebel

**Author notes:** denotes equal contributions (Huahua Sun, Feng Liu).

## Abstract

Anopheles mosquitoes are the sole vectors of malaria and other diseases that represent significant threats to global public health. While adult female mosquitoes are responsible for disease transmission, the pre-adult larval stages of the malaria vector *Anopheles coluzzii* and other mosquitoes rely on a broad spectrum of sensory cues to navigate their aquatic habitats efficiently to avoid predators and search for food. Of these, mosquito larvae rely heavily on volatile chemical signals that directly activate their olfactory apparatus. Because most studies on mosquito olfaction focus on adults, a paucity of attention has been given to the larval olfactory system, in which the peripheral components are associated with the sensory cone of the larval antennae. To address this, we have investigated the electrophysiological response profile of the larval sensory cone in Anopheles mosquitoes. We found that the larval sensory cone is particularly tuned to alcohols, thiazoles and heterocyclics. Furthermore, these responses can be assigned to discrete groups of sensory cone neurons with distinctive, dose-dependent odorant-response profiles that also provide larvae with the ability to discriminate among compounds with similar chemical structures. A correlation analysis was conducted to determine the relationship between specific larval chemosensory receptors and the response profiles of sensory cone neuron groups. These studies reveal that the larval sensory cone is a highly sophisticated organ that is sensitive to a broad range of compounds and is capable of a remarkable degree of chemical discrimination. Taken together, this study presents critical insights into olfactory coding processes in *An. coluzzii* larvae that further our understanding of larval chemical ecology and will contribute to the development of novel larval-based strategies and tools for mosquito control and the reduction of vector-borne disease transmission.

## Introduction

While significant progress has been made advancing our understanding of the molecular and cellular basis of the olfactory system of adult mosquitoes (Montell and Zwiebel, 2016; Lutz et al., 2017; Zwiebel and Takken, 2004), considerably less is known about olfactory processes during the mosquito’s pre-adult larval and pupal life-stages where, paradoxically, the majority of successful malaria control strategies have been historically focused (Floore, 2006; Tusting et al., 2013). Anopheline larvae develop aquatically across four stages, known as instars, for approximately 10days, depending on species and ambient temperature (Clements, 2000). Larval populations exist in restricted and often tenuous habitats where they compete with each other as well as a range of other life forms, some of which are predatory, for food and survival. Rather than simply representing an immature life stage with relatively narrow perspectives, larval-stage mosquitoes are in fact highly complex, independent organisms that survive only by employing a wide range of physiological and sensory systems, with many of the same components and characteristics of similarly tasked adult processes.

Fundamental behaviors such as the acquisition of nutrients and avoidance of danger rely on the larval olfactory system to detect and respond to a complex set of chemical cues marking potential food sources (Xia et al., 2008), predators (Sih, 1986) and other aspects of their aquatic environments. Unlike Aedine or Culicine larvae, which dive to the bottom of their aquatic habitats to food search, Anopheline larvae remain parallel to the surface, using their mouth brushes to filter potential food sources, such as microorganisms and other organic compounds (Fillinger et al., 2004; Gimnig et al., 2001). The surface water contains relatively high amounts of nutrients, particulate and dissolved organic material, and various microorganisms, which emit volatile and non-volatile chemical cues that attract Anopheline larvae (Danos et al., 1983; Wotton et al., 1997).

Mosquitoes and nearly all dipteran larvae share an array of homologous anterior appendages that comprise their principal chemosensory organs. The larval antennae extend distally from the head and include, at the apex, both a sensory cone and a peg organ which are respectively considered olfactory and gustatory organs (Lutz et al., 2017; Nicastro et al., 1998; Xia et al., 2008; Zacharuk et al., 1971). In *D. melanogaster*, the larvae olfactory apparatus consists of two bilaterally symmetrical dorsal organs, in which 21 odorant receptor neurons (ORNs) expressing 23 odorant receptors (ORs) as well as Orco, the obligate co-receptor, have been identified (Kreher et al., 2005). In the dengue virus vector mosquito *Aedes aegypti*, the larval sensory cone is innervated by 12-13 typical bipolar neurons expressing 24 *Or* genes, 15 of which are larvae specific (Bohbot et al., 2007; Zacharuk et al., 1971). In *An. coluzzii* and *An. gambiae* (previously known as the ‘M’ and ‘S’ forms of the *An. gambiae* species complex (Coetzee et al., 2013), the larval antenna contains 12 distinct ORNs, in which an equal number of ORs (four of which are larvae specific) are co-expressed along with the *Anopheles orco* gene (Liu et al., 2010; Xia et al., 2008). In addition, at least four *An. coluzzii* variant ionotropic receptors (IRs) are expressed on IR chemosensory neurons (IRNs) within the larval antennae, supporting the hypothesis that ORN- and IRN-based chemosensory signaling represent two distinct pathways involved in olfactory-driven behaviors of Anopheline larvae (Liu et al., 2010; Xia et al., 2008). These studies demonstrated that *An. coluzzii* larvae depend on their antennae and *orco* expression to respond behaviorally to a range of natural and synthetic odorants, such as organic decay cresol derivatives (2-methylphenol, 3-methylphenol, 4-methylcyclohexanol) and the insect repellent DEET (Liu et al., 2010; Xia et al., 2008). Similarly, siRNA-mediated gene silencing of the *An. gambiae Ir76b* co-receptor specifically altered *Anopheles* larval responses to butylamine (Liu et al., 2010). Functional characterization of several larval *Anopheles* ORs via heterologous expression in *Xenopus* oocytes shows that larval ORs respond to a range of odorants that have significant roles in larval chemical ecology (Xia et al., 2008).

Thus far, the characterization of *in vivo* electrophysiological responses of mosquito ORNs has been exclusively restricted to adult stages, with a limited number of studies restricted to terrestrial larvae in other insects. Extracellular electrophysiological recordings from the dorsal organ in *Drosophila* revealed background firing from multiple ORNs with varying amplitudes, which proved difficult to fully sort but nevertheless uncovered several odorants that elicit excitatory neuronal responses, including 2-methylphenol, acetophenone, and benzaldehyde (Kreher et al., 2005). In addition, recordings from the caterpillar stages of *Spodoptera littoralis* moths revealed that its olfactory sensilla were sensitive to sex pheromone components and plant odors (Poivet et al., 2012; Rharrabe et al., 2014). In contrast, *in vivo* electrophysiological characterizations of larval olfactory responses have thus far not been reported in any mosquito, most likely as a result of the inherent challenges of the larval aquatic environment. This paucity of attention is paradoxical to the importance of mosquitoes as global disease vectors as well as the central role that larval stages play in both the mosquito life cycle and historically effective control strategies. To address this knowledge gap, we have systematically characterized the olfactory responses of the *An. coluzzii* larval sensory cone using electrophysiological approaches adapted from adult single sensillum recording (SSR). Taken together, these studies demonstrate that, as is the case for Anopheline adults, larvae have a complex and multi-faceted peripheral olfactory apparatus that provides a significant degree of chemosensory discrimination that is specifically adapted for the requirements of larval life.

## Materials and Methods

### Mosquito Rearing

*An. coluzzii* adults were reared at 27°C, 75% humidity under a 12h light/12h dark photoperiod and supplied with 10% sucrose water in the Vanderbilt University Insectary (Fox et al., 2001; Suh et al., 2016). For stock propagation, 5- to 7-day-old mated females were blood fed for 30-45min using a membrane feeding system (Hemotek, Lancaster, UK) filled with defibrinated sheep blood purchased from Hemostat Laboratories (Dixon, CA, USA).

Mosquito larvae were reared in distilled water at 27°C under the standard 12h light/12h dark cycle, with approximately 300 larvae per rearing pan in 1L H_2_O. The larval food was made from 0.12g/mL Kaytee Koi’s Choice premium fish food (Chilton, WI, US) plus 0.06g/mL yeast in distilled water and subsequently incubated at 4°C overnight for fermentation. For 1^st^ and 2^nd^ instar larvae, 0.08mL larval food was added into the water every 24h. For 3^rd^ and 4^th^ instar larvae, 1mL larval food was added. Only 3^rd^ instar larvae were used in the experiments.

### Larval Electrophysiology

Electrophysiological recordings were conducted on the *An. coluzzii* larval antennal sensory cone using an adaptation of the well-established single sensillum recording technique (SSR) (Liu et al., 2013). Here, 3^rd^ instar larva were mounted on 76×26-mm microscope slides (Ghaninia et al., 2007), and using double-sided tape the larval antennae were fixed to a cover slip, resting on a small bead of dental wax to facilitate manipulation with the cover slip placed at approximately 30 degrees relative to the head. Once mounted, the specimen was positioned onto an Olympus BX51WI microscope, and antennae were viewed at high magnification (1000×). Tungsten microelectrodes (sharpened in 10% KNO_2_ at 10V) were connected to a pre-amplifier (Syntech universal AC/DC 10x, Syntech, Hilversum, The Netherlands) and used for both the grounded reference electrode that was inserted into the larval compound eye using a micromanipulator (Olympus BX51W1) and the recording electrode inserted into the base of the antennal sensory cone to complete the electrical circuit to record olfactory sensory neuron (OSN) potentials extracellularly (Den Otter et al., 1980). Controlled manipulation of the recording electrode was performed using a Burleigh micromanipulator (Model PCS6000). The pre-amplifier was connected to an analog-to-digital signal converter (IDAC-4, Syntech, Hilversum, The Netherlands) at a sample rate of 96,000 samples per second and 10× amplification, which in turn was connected to a PC for signal recording and offline analysis.

### Odorant preparation and stimulus application

Compounds with highest purity, typically ≥99% (Sigma-Aldrich), were diluted in paraffin oil or dimethyl sulfoxide (DMSO) or Diethyl Pyrocarbonate (DEPC)-treated ddH_2_O to make v/v (for liquids) or m/v (for solids) solutions at certain concentrations. For each compound, a 10-μL portion was dispersed onto filter paper (3×10mm), which was then inserted into a Pasteur pipette to create the stimulus cartridge. A sample containing the solvent alone served as the control. The airflow across the antennae was maintained at a constant 20 mL/s throughout the experiment. Purified and humidified air was delivered to the preparation through a glass tube (10-mm inner diameter) perforated by a small hole 10cm away from the end of the tube into which the tip of the Pasteur pipette could be inserted. The stimulus was delivered to the larval, antennal sensory cone by inserting the tip of the stimulus cartridge into this hole and diverting a portion of the air stream (0.5 L/min) to flow through the stimulus cartridge for 500ms (0.5 s) using a Syntech stimulus controller CS-55 (Syntech, Hilversum, The Netherlands). The distance between the end of the glass tube and the antennae was ≤1 cm. For dose-response relationships, odorants were presented in increasing doses. Signals were recorded for 10 s, starting 1 s before stimulation, and the background action potentials from the entire sensory cone were counted off-line over one 0.5-s period before the stimulus and over four 0.5-s periods during and after stimulation. Spike rates observed during the 0.5-s stimulation/post-stimulation windows were normalized by subtracting the pre-stimulus (background) activities observed in the preceding 0.5 s, with counts recorded in units of spikes/s. Lastly, odorant responses, described as Δ spikes/s, were normalized by subtracting the solvent responses in each individual recording.

### Spike sorting

The action potential recordings from larval sensory cones were initially transformed into spikes and sorted, using AutoSpike (Syntech, Hilversum, The Netherlands), into three discrete groupings based on the amplitude histograms as follows: (1) group “A” neurons with large spike amplitudes >800μV; (2) group “B” neurons with medium spike amplitudes between 400 and 800μV; and (3) group “C” neurons with small spike amplitudes ≤400μV. Each recording was initially validated for spike sorting based on the group C neuron amplitude threshold in the pre-stimulus (background) conditions, such that all group C neuron spikes in this interval must have an amplitude ≥200 μV. If this criterion was not met, the recording was not used for sorting analysis. Spikes were counted across a single 0.5-s pre-stimulus interval as well as four 0.5-s (2 s total) post-stimulus measurements. “Pre-stimulus” is defined as the time immediately preceding odorant delivery and “Post-stimulus” as the time immediately following the onset of neuronal activation. Neuronal responses (Δ spikes/0.5 s) were calculated by subtracting the pre-stimulus activity from the spike number observed during each 0.5-s post-stimulus measurement and further corrected by subtracting solvent-alone response for each recording.

### Odorant Response/Or Functionality Correlation Analysis

Correlative analyses between larval sensory cone neuron group responses and the functional profiles of larvally expressed *An. coluzzii* ORs 1, 2, 6, 10, and 48 were carried out using the Drosophila empty neuron system dataset (Carey et al., 2010) and for ORs 28, 37, 40 using a *Xenopus* expression system dataset (Wang et al., 2010; Xia et al., 2008). Here, responses to odorants used in both those OR functional studies and *in vivo* electrophysiological recording (this study) were selected and normalized to the largest response in each corresponding assay. The normalized odorant response profiles of each functionally characterized larval OR were graphed along the x-axis, and the normalized odorant response profiles of each of the three ABC neuron groups characterized in the larval sensory cone were graphed along the y-axis. A Pearson’s correlation analysis incorporating a linear regression between these data sets was performed using Prism v6 (GraphPad Software, San Diego, CA). The correlation coefficient parameter, R2, was generated to assess how strong a relationship existed between the two variable datasets. *F tests* with a p<0.05 were considered a significant correlation.

## Results

### Broad response profiles of the larval sensory cone of *An. coluzzii*

The larval antennae of mosquitoes encompass several orthologous structures that are morphologically conserved across individuals and instars of larvae. One of these structures, the sensory cone, is the site of peripheral chemosensory signal transduction for olfaction (Zacharuk et al., 1971). While previous studies have cataloged and functionally analyzed the OR repertoire of the larval antenna *in vitro* as well as the role of ORs and IRs in larval behavioral responses (Liu et al., 2010; Xia et al., 2008), we now report a comprehensive characterization of *in vivo* peripheral neuronal response profiles from the larval sensory cone in *An. coluzzii*. These results provide a direct link between those responses and corresponding larval behaviors, as well as further illustrating the inherent complexity of the odor-coding paradigms of mosquito larvae. Initial electrophysiological surveys of the sensory cone on the tip of larval antennae consistently revealed the presence of a multifaceted neuronal background activity. Stimulus-independent action potential spikes from most recordings could be manually and computationally separated into several discrete groups based on their amplitudes (Figure S1A and S1B), which indicates the presence of multiple neurons within an individual antennal sensory cone.

In order to investigate the broad olfactory profiles of *An. coluzzii* larvae, we initially examined collective (unsorted) neuronal responses after challenging the sensory cone with an in-house panel of 281 odorants, spanning 12 chemical classes and including aromatics, heterocyclics, alcohols, ketones, aldehydes, thiazoles, sulfurs, terpenoids, carboxylic acids, amines, esters and others. Several of these compounds are components of larval food sources, predator emanations, and oviposition site volatiles (Figure 1A, Supplemental Table S1) that have been shown previously to evoke behavioral activity in *An. coluzzii* larvae (Xia et al., 2008). Surprisingly, less than 20% of the 279 unitary odorants screened at 10^−2^ dilution elicited very strong olfactory responses (defined as ≥80 Δ spikes/s). Indeed, the frequency of robust (defined as 40-80 Δ spikes/s), modest (defined as 10^−40^ Δ spikes/s) and low responses (defined as <10 Δ spikes/s) was 29.1%, 41.6% and 11.7%, respectively (Figure 1A, Supplemental Table S2). Interestingly, the frequency of very strong excitatory responses was significantly higher among alcohols (67.9%), thiazoles (62.5%) and heterocyclics (38.1%) than terpenoids (4.8%), esters (2.3%) and carboxylic acids (0%). A collective odorant-tuning curve spanning larval responses to all 281 odorants revealed a broad distribution with a very small kurtosis value (k=0.136), which measures the peakedness of the distribution as a reflection of its specialization (Figure 1B) (Carey et al., 2010; Wetzel et al., 1999). Within the entire larval sensory cone tuning curve, it is very clear that alcohols, thiazoles and heterocyclics with the strongest responses are clustered in the center, ketones with the modest response in the middle part of either side, while acids with the weakest response are placed at each end (Figure 1B). As expected, the apex responses are derived from three alcohols (1-pentanol, trans-3-hexen-1-ol and 1 butanol) (Figure 1B). Taken together, these data suggest that the larval sensory cone is a generalist sensory appendage that displays a modest coding bias tuned toward alcohols, thiazoles and, to a somewhat lesser extent, heterocyclic-based cues.

**Figure 1.**
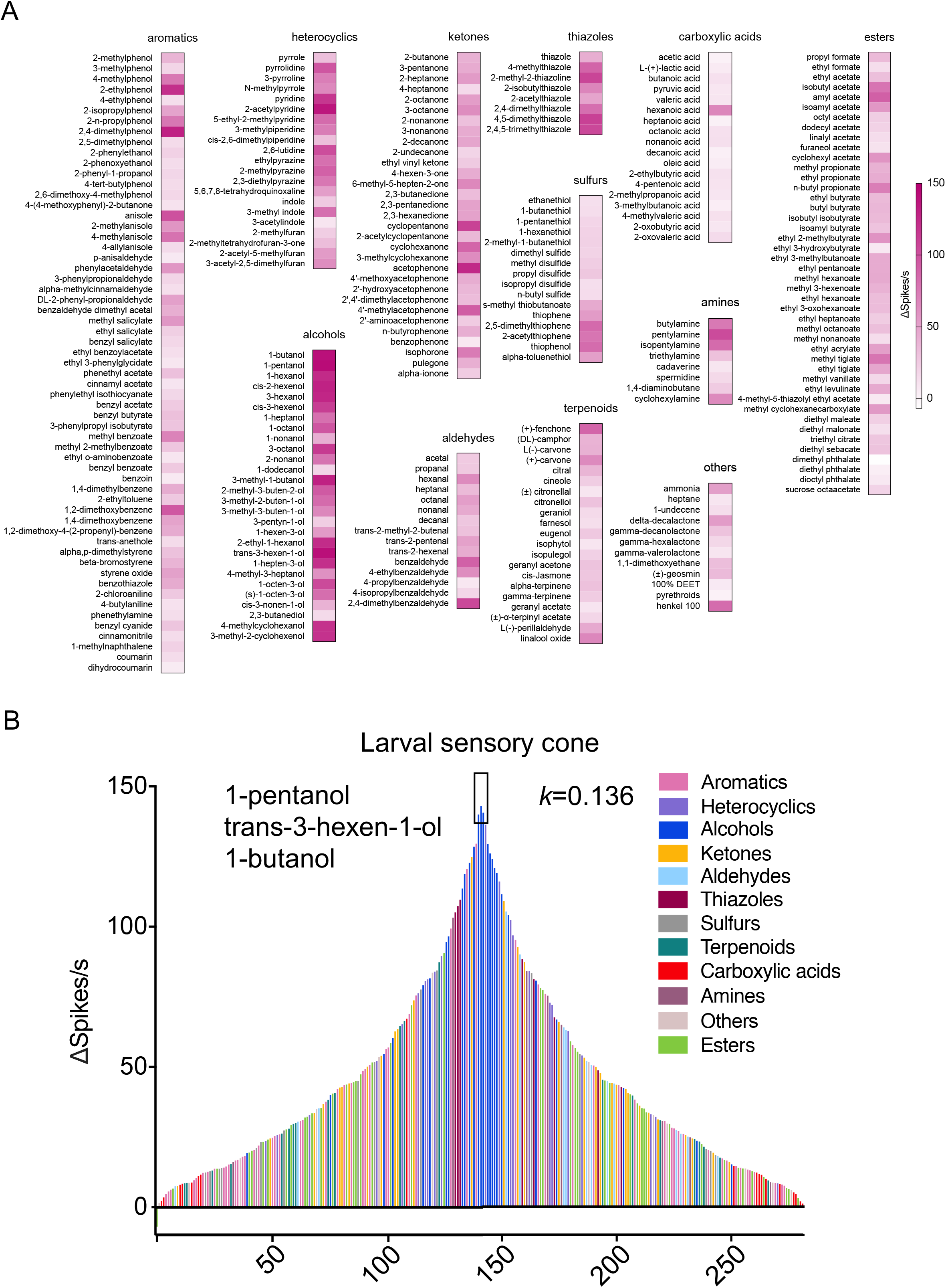
(A) Response profile of the larval, antennal sensory cone to 281 compounds of diverse chemical groups (n=6-10). (B) Tuning curve reveals the breadth of the larval, antennal sensory cone. The 281 compounds are distributed along the x-axis according to the strengths of the responses they elicited from the sensory cone. The odors that elicited the strongest responses are near the center of the distribution; those that elicited the weakest responses are near the edges. Negative values indicate inhibitory responses. Each odorant is color coded based on their chemical classes. The kurtosis (k) value, as a statistical measure of ‘peakedness’, is shown on the right side of the plot.

### Odorant tuning in the larval sensory cone is concentration dependent

Our screening revealed a subset of 27 odorants that elicited strong responses across one or more neuron groups. Within those, we next addressed whether the sensory cone displayed dosedependent responses, especially at low concentrations that are more likely to represent the natural environment of mosquito larvae. *An. coluzzii* larval sensory cone neurons were therefore interrogated using the robust response subpanel of 27 odorants at concentrations spanning five orders of magnitude (10^−5^–10^−1^ dilution; Figure 2A, C, Supplemental Table S3). Not surprisingly, none of the 27 odorants elicited more than a modest response at low concentration (10^−4^ and 10^−5^ dilution). Furthermore, of those that did elicit a robust (≥40 Δ spike/s) response at 10^−2^ dilution, only 13 maintained this intensity at lower (10^−3^) dilutions (Figure 2A, Supplemental Table S3). A tuning curve analysis illustrates these odorants evoke broad, concentration-dependent responses with characteristically low peakedness and are modestly biased toward 2,4,5-trimethylthiazole and 4-methylcyclohexanol at lowest concentrations (10^−5^ and 10^−4^ dilutions, respectively), acetophenone at middle concentrations (10^−2^ and 10^−3^ dilutions), and butylamine at the highest concentration (10^−1^ dilution; Figure 2B). Across this range, sigmoidal dose–response curves showing clear saturation at 10^−2^ dilutions were only observed for two aromatics (2-ethylphenol, 2,4-dimethylphenol) and two ketones (cyclopentanone, acetophenone) (Figure 2C), further illustrating the larval sensory cone’s response complexity.

**Figure 2.**
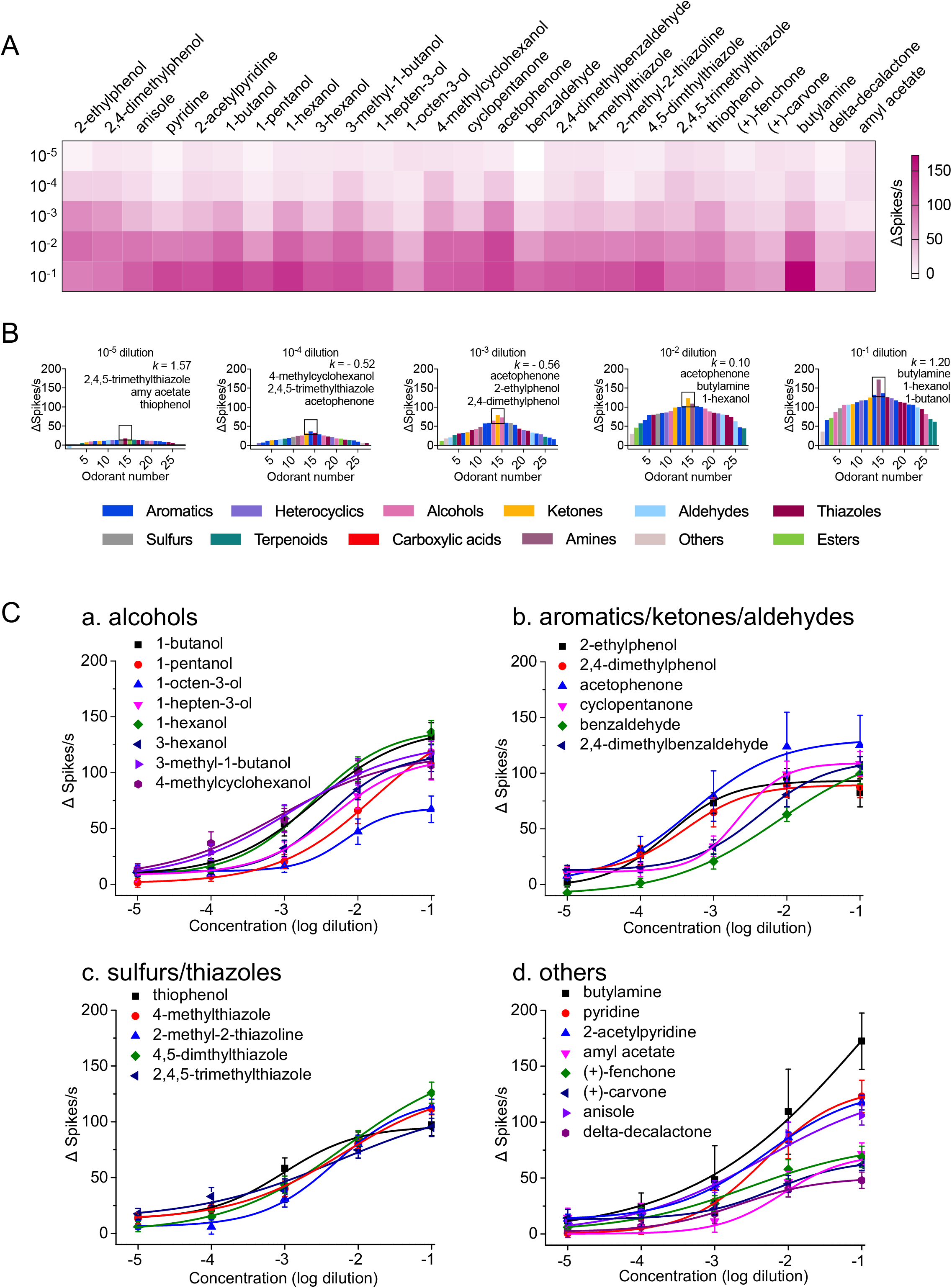
(A) Dose-dependent response to a panel of 27 odorants displayed along heat map quadrants. (B) Tuning curves of the larval antennal sensory cone to odorant dilutions across five orders of magnitude. The 27 compounds are distributed along the x-axis according to the strength of the responses they elicited from the sensory cone. The odorants that elicited the strongest responses are near the center of the distribution; those that elicited the weakest responses are near the edges. The three odorants eliciting the highest firing frequency in the sensory cone are noted for each dilution: (C) Dose–response curves of the larval antennal sensory cone to these 27 odorant stimuli. n=6-10 for each odorant dilution in each compound.

### Fine-scale temporal dynamics of neuronal responses

Although these recordings do not facilitate an unambiguous molecular mapping of these chemosensory neurons into, for example, IRN or ORN categories, we nevertheless elected to sort responses into three broad neuronal groups based on their spike amplitude. These are the “A group” with large amplitude spikes (>800μV), “B group” with medium amplitude spikes (400-800μV), and “C group” with small spikes (≤400μV; Figure S1A/B). As detailed below, while some odorants display selective responses, a wide variety of odorants, including 4-methylphenol and most of the alcohols, activated and increased the firing frequency of multiple groups of neurons on the larval sensory cone of *An. coluzzii* (Figure S1C).

While this approach is restricted to the 128 odorants that evoke supra-threshold responses and does not provide single neuron-level discrimination, it does allow us to distinguish three discrete classes of larval chemosensory neurons. Furthermore, in order to examine the temporal dynamics of odorant responses across the larval sensory cone, firing frequencies could be quantified for each neuron group across four 0.5-s intervals (a total of 2s) during and after stimulation with odorant delivery restricted to the first 0.5s window (Figure 3, bold, red bar; Supplemental Table S4). Not surprisingly, sustained tonic stimulation across all A, B, and C neuron groups for the entire 2s was elicited only by the complex Henkel 100 odorant mixture (Henkel AG, Dusseldorf, Germany; Figure 3, red arrowhead), while several unitary compounds were able to elicit prolonged 2-s responses within specific groups of sensory cone neurons (Figure 3, Supplemental Table S4). Of these, only three aromatics (2-ethylphenol, 2-isopropylphenol 1,2-dimethoxy-4-(2-propenyl)-benzene) and one heterocyclic (3-methylindole) evoked sustained excitatory responses solely across the A-group neurons. However, across B-group neurons, selective and sustained excitatory responses were evoked by a handful of compounds comprising four aromatics (2-n-propylphenol, 2,4-dimethylphenol, methyl salicylate, benzothiazole), two heterocyclics (pyrrolidine, 3-pyrroline), two alcohols, (1-nonanol, 2-ethyl-1-hexanol), two ketones (2-decanone, 4’-methoxyacetophenone) and a single aldehyde (2,4-dimethylbenzaldehyde) together with two terpenoids ((+)-carvone and L-(-)-perillaldehyde). With 1-heptanol, 1-octanol, hexanoic acid and 2,3-butanedione as the exceptions, C-group neurons did not display tonic excitation in response to our odorant panel (Figure 3, Supplemental Table S4).

**Figure 3.**
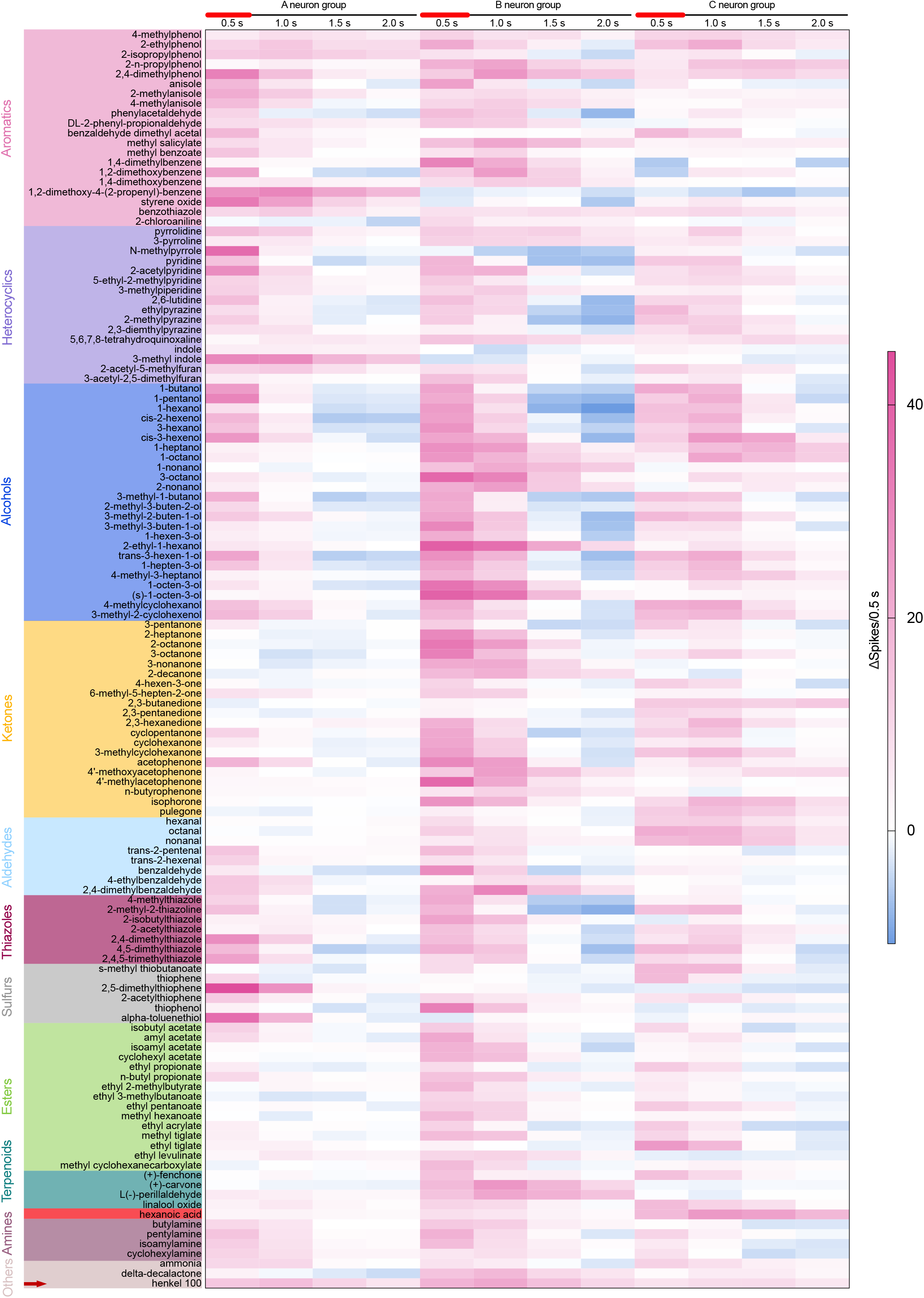
Response profile of three neuron groups to 128 odors eliciting ≥40 spikes in the sensory cone (n=6-10). Four 0.5-s intervals of neuron firings were quantified after the initiation of responses. The odorant puff was delivered in the first 0.5s, as indicated by red and bold line.

Beyond these sustained responses, some odors, including 1,2-dimethoxy-4-(2-propenyl) benzene, styrene oxide, 2,5-dimethylthiophene, 3-methylindole and alpha-toluenethiol, elicited very strong selective responses solely in A-group neurons (Figure 3, Supplemental Table S4). A major portion of ketones and esters activated both B- and C-group neurons but not A neurons; several heterocyclics uniformly evoked moderate responses from all three types of neurons; but most alcohols predominantly elicited responses from only B-group neurons. We also observed that most aliphatic alcohols elicited an initial 1-s excitatory response followed by a similarly timed inhibitory response across both the A- and B-group sensory cone neurons, while esters were more likely to show this response pattern in C-group sensory cone neurons (Figure 3, Supplemental Table S4).

In addition to this phenomenon, phasic and tonic response patterns were sometimes present in larval sensory cone neuron firing. Here, the responses of A-group neurons to N-methylpyrrole, 1-pentanol and 2,5-dimethylthiophene, and B-group neurons to thiophenol were of a very typical phasic firing pattern. In contrast, the responses of A-group neurons to 3-methylindole and 1,2-dimethoxy-4-(2-propenyl)-benzene, B-group neurons to methyl salicylate, 2-ethyl-1-hexanol, (s)-1-octen-3-ol, L-(-)-perillaldehyde and (+)-carvone, and C-group neurons to hexanoic acid and 2,3-butanedione were of a more tonic mode, with super-sustained firing (Figure 3, Supplemental Table S4).

We also conducted a more detailed examination using a narrower subset of 20 compounds by further dissecting the 0.5-s windows into 0.1-s quantitative intervals to better understand the temporal dynamics of sensory neuronal responses (Figure 4). This analysis revealed that, while many odorants seemed to activate only A-group (1,2-dimethoxy-4-(2-propenyl)-benzene, 3-methylindole, 2,5-dimethylthiophene, N-methylpyrrole) neurons or B-group (1,4-dimethylbenzene, 2-ethyl-1-hexanol, (s)-1-octen-3-ol, thiophenol, (+)-carvone) neurons, restrictive C-group excitation was observed for hexanoic acid and 2,3-butanedione stimulation. Indeed, these data reveal significant complexity in these responses, including potential inhibitory interactions among these three groups of sensory cone neurons in which the strong activation of one neuron group inhibits the firing of other neuron group(s) resulting in latent excitatory responses. For example, the strong excitation of A-group neurons by 2,4-dimethylphenol seemed to temporally inhibit responses from B- and C-group neurons, resulting in a delayed and potentially weakened response (Figure 4J). Similarly, peak excitation of B-group neurons by 2,4-dimethylbenzaldehyde was only observed when A-group excitation diminished (Figure 4K). Moreover, hexanoic acid elicited prolonged responses, initially consisting of moderate responses across all larval cone neuron groups in the 1^st^ 0.5s, followed by increasingly stronger C-group neuron peak responses in the 2^nd^–4^th^ 0.5-s (post-odorant) intervals (Figure 4T).

**Figure 4.**
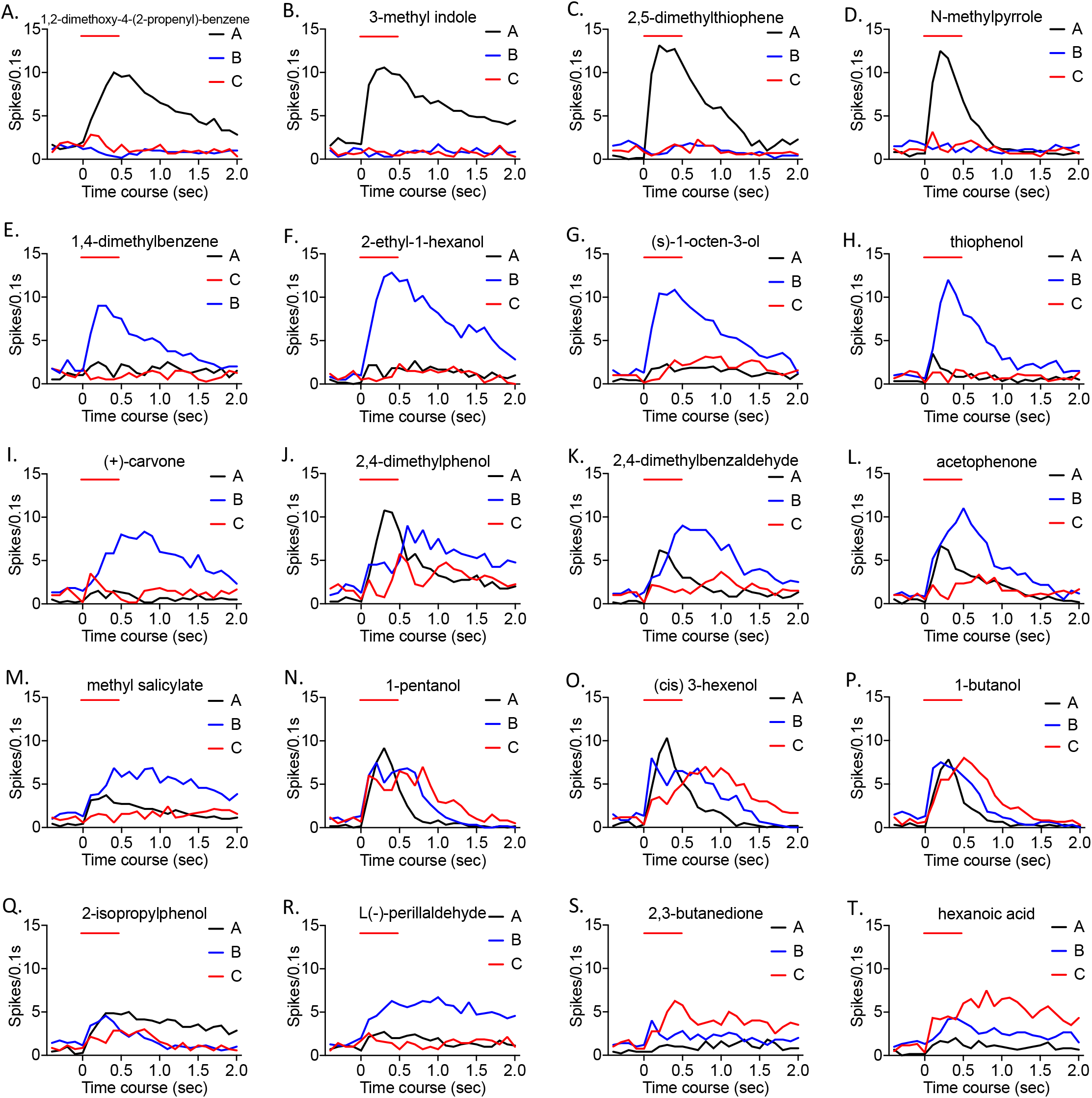
Temporal dynamics of responses of different neuron groups to 20 highly responsive odorants (n=6-10). Every 0.1-s interval of neuron firing was quantified for 0.5s before and 2s after odorant stimulation. Temporal responses from A-group neurons (black lines), B-group neurons (blue lines) and C-group neurons (red lines) are displayed individually for each stimulus.

When examining neurons with a firing rate ≥40 spikes/s, which generally corresponds to 30% of the maximum response, we found that during odorant delivery (the initial 0.5-s window), B- and C-group neurons displayed broader tuning curves, with kurtosis values of 0.03 and −0.68, respectively, than A-group neurons, with a kurtosis value of 1.32 (Figure S2A), perhaps indicating a degree of specialization. Furthermore, during the odorant delivery window, the compounds eliciting peak responses in the A-, B-, and C-group neurons are distinct from each other as well as from those when collectively examining the larval sensory cone (Figure S2). For example, the most intense responses evoked from A-group neurons are generated by 2,5-dimethylthiophene, N-methylpyrrole and alpha-toluenethiol (Figure S2), which does not overlap with the collective responses across the larval sensory cone (Figure 1B).

Additional complexity was observed when these dose-dependent, larval sensory cone responses were further sorted into spike amplitude-dependent neuronal groups and temporally examined (Figure S3, Supplemental Table S5). These data show that sustained tonic stimulation across the entire 2-s recording paradigm was broadly correlated but, importantly, not entirely restricted to the highest stimulus concentrations. Indeed, two aromatic odorants (2,4-dimethylphenol and 2-ethylphenol) elicited sustained 2-s activation of A- and B-group neurons at dilutions as low as 10^−4^. If one examines the less stringent parameter of activation for only the initial 1-s stimulation window, several odors were capable of activating specific neuron groups at very low concentrations, very rarely at 10^−4^ dilutions (e.g. 4-methylcyclohexanol and acetophenone for B-group neurons and 2-methyl-2-thiazoline for C-group neurons) and beyond that became more commonplace with increasing odorant concentration. At the opposite extreme, some odorants only activated specific neurons at the highest concentration (e.g. pyridine only evoked responses in A-group neurons at 10^−1^ (Figure S3). These disparities in dose-dependent response profiles indicate the variance in potency of compounds to activate different neuron groups in the sensory cone. Taken together, these findings indicate that different groups of neurons in the larval sensory cone use a variety of approaches to cover distinctive odor spaces in the aquatic *An. coluzzii* larvae.

### Diverse response patterns of the larval sensory cone to structurally similar odorants

We next sought to determine how mosquito larvae might discriminate among odorants with similar chemical structures. While our initial expectation was that odorants with conserved structures would elicit similar responses across the larval sensory cone, as was observed for two alcohols (1-butanol and 1-pentanol) and several anisoles (anisole, 2-methylanisole and 4-methylanisole) (Figure S4A, B), striking differences were noted for other structurally similar stimuli. For example, the strength of responses to carboxylic acids appeared to be highly dependent on a discrete six-carbon-chain length. Robust responses from C-group neurons were elicited by hexanoic acid, but very weak responses were observed with shorter and longer chain acids (Figure 5A). Additionally, different neuron groups in the sensory cone strongly responded to multiple aromatic sulfurs, but none of the aliphatic sulfurs (Figure 5B). The location of alkane groups on a benzene ring may be another indicative factor, as responses between 2-ethylphenol and 4-ethylphenol, and between 2,4-dimethylphenol and 2,5-dimethylphenol differed dramatically, especially in the B-group neurons (Figure 5C). Furthermore, methyl groups on a thiazole ring demonstrated a higher response strength than thiazole rings alone (Figure 5D). In addition, we observed considerable variance in response to similar compounds among different neuronal groups. While several cresols and cyclic ketones with similar structures elicited aligned response in the C-group neurons, the response of A- and B-group neurons to these compounds were much more complex (Figure S5C and D). For instance, 2-isopropylphenol evoked a significantly larger response in A neurons than 2-n-propylphenol. However, the opposite response was observed in the B-group neurons. Taken together, these results indicate that the neurophysiological responses of *An. coluzzii* larvae display a robust ability for detecting and discriminating complex chemical components from their aquatic habitats, despite manifesting only a small fraction of the neuronal and molecular repertoire present in the adult olfactory appendages.

**Figure 5.**
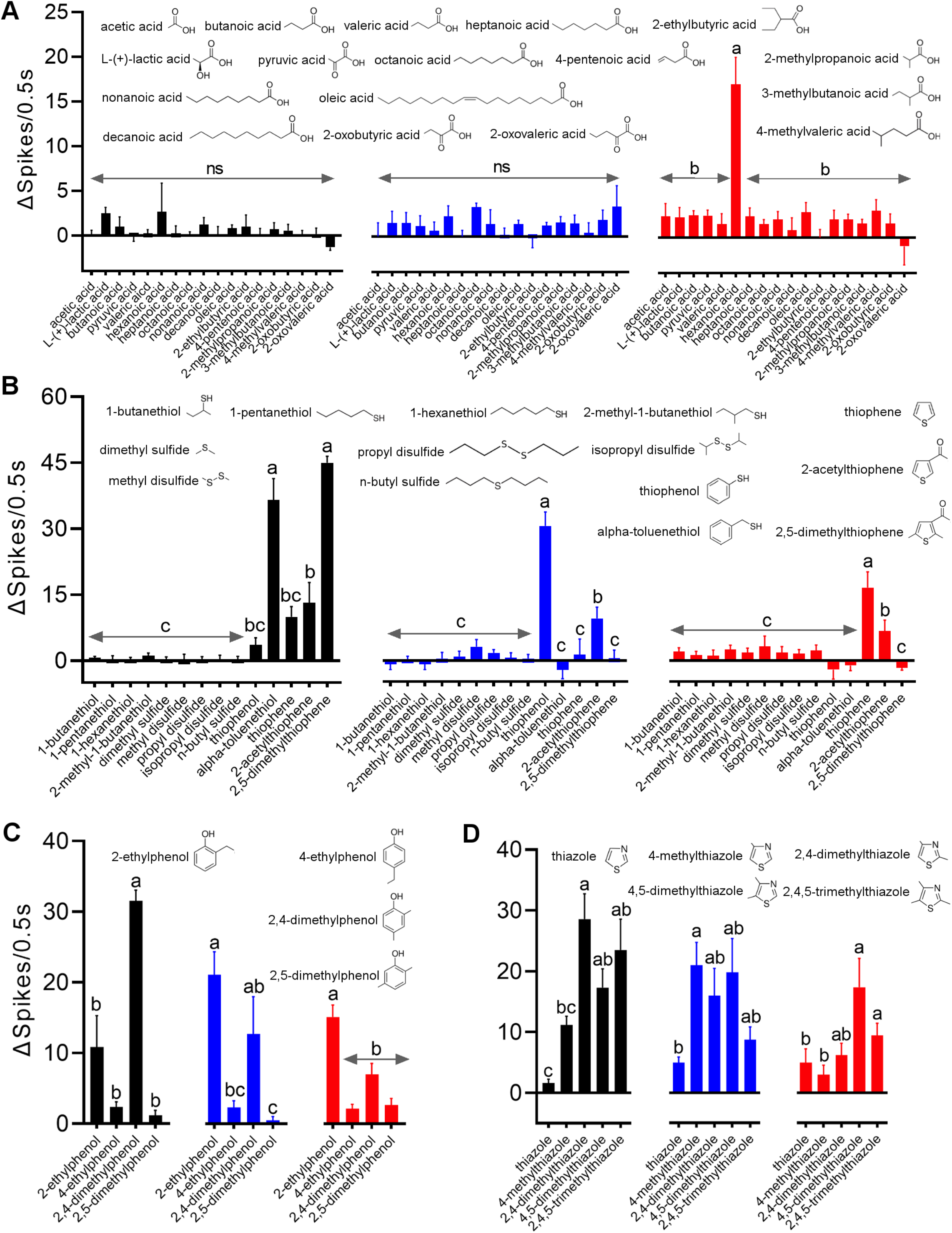
Responses of three larval cone neuron groups to odorants with similar structures (n=6-10). Neuronal responses are color coded: A-group neurons (black), B-group neurons (blue) and C-group neurons (red). (A) Response of sensory cone neurons to aliphatic carboxylic acids shows hexanoic acid selectively evoking a significant C-group response when compared with other acids. Differential responses to (B) aliphatic and aromatic sulfurs; (C) ethylphenol-derived compounds; and (D) thiazole-derived compounds with increased alkane branches in the heterocyclic ring. One-way ANOVA Tukey test was applied in the statistical analysis, with p<0.05 indicating significant difference.

### Correlations between neuronal and OR responses

The functional characteristics of specific chemosensory receptors (e.g. ORs and IRs) expressed in the dendritic membrane of mosquito ORNs and IRNs housed in adult olfactory sensilla and larval sensory cone largely determine the response profiles of those neurons. Previous studies have functionally characterized and examined the odor coding of eight larval ORs in “M-form” *An. gambiae* (Carey et al., 2010; Wang et al., 2010; Xia et al., 2008), now renamed *An. coluzzii* (Coetzee et al., 2013).These studies make it possible to correlate the functionality of those larval ORs with the activity of peripheral neuron groups in the larval sensory cone and, thereby, uncover the potential role of specific *Or* genes in each neuron group. Specifically, of those eight larval ORs, a significant correlation was observed between the response profiles of C-group neurons and that of OR40, which gives a correlation coefficient parameter R2 =0.417, suggesting that *Or40* is likely to be expressed in at least one of the ORNs that make up the larval sensory cone C-group neurons (Figure 6U). In addition, we also found a significant correlation between the response profiles of OR28 with A-group neurons and OR48 with B neurons, with R2 values of 0.314 and 0.390, respectively, suggesting that *Or28* is likely to be expressed in one of the A-group neurons and *Or48* in one of the B-group neurons (Figure 6 M, W). None of the other correlations in the OR/neuronal response profiles was significant, leaving us unable to speculate as to their distribution across larval sensory cone neuron groups.

**Figure 6.**
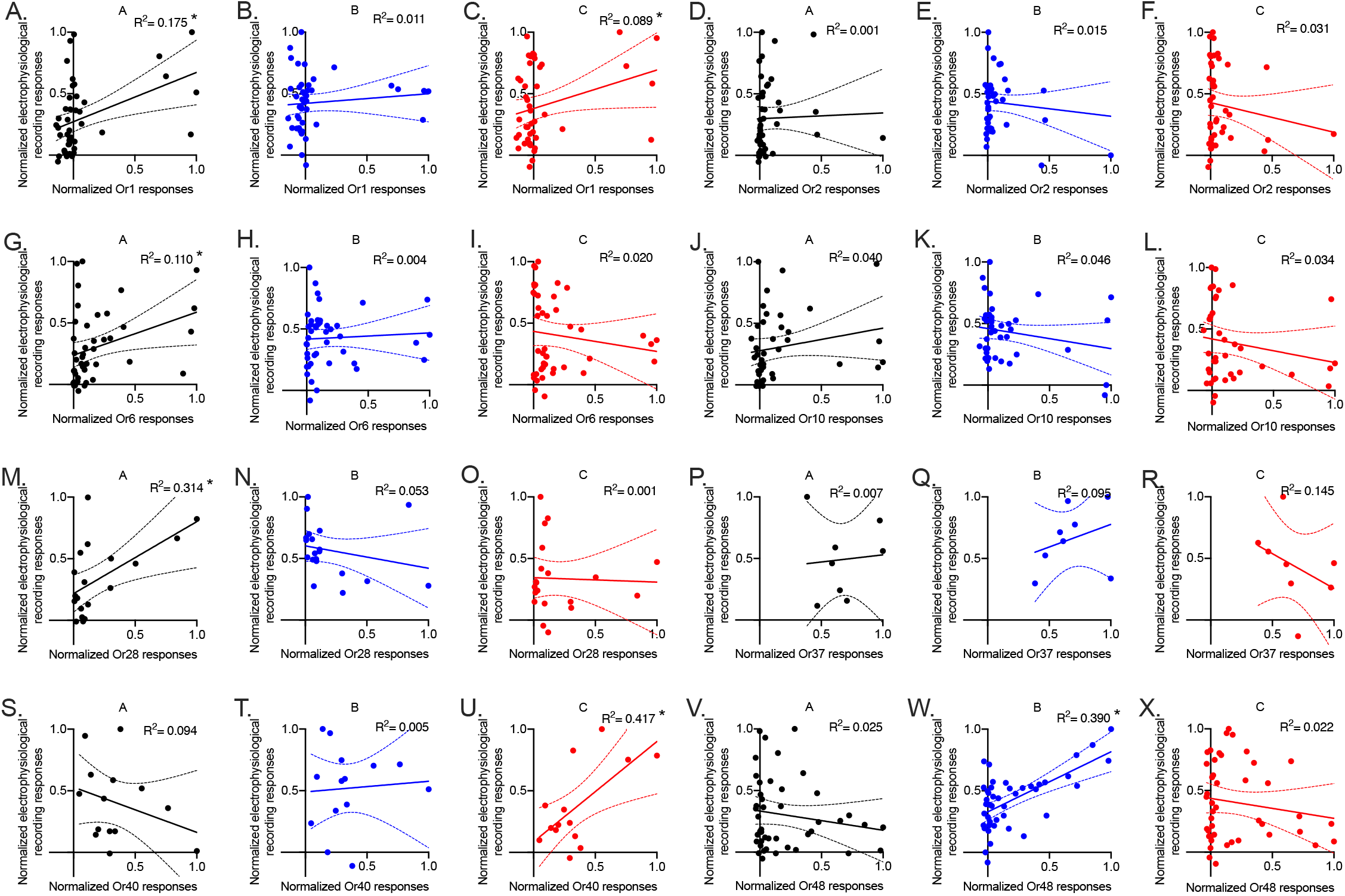
Correlative analysis of response profiles of different neuron groups and heterologously expressed *An. coluzzii* ORs. The response profiles of ORs1, 2, 6, 10, and 48 were retrieved from Carey et al. (2010) and the response profiles of ORs28, 37, and 40 were retrieved from Wang et al. (2010). Responses to the overlapping compounds in both OR functional studies and *in vivo* electrophysiological recordings in this study were first normalized to the corresponding largest response, and then correlation analysis was conducted. *F* tests with a p value less than 0.05 are considered as significant correlations.

## Discussion

Larval Anophelines are essentially aquatic insects with a unique chemical ecology. As is the case for the adult stages, mosquito larvae utilize their olfactory system together with other sensory modalities to respond to the presence of predators as well as identify and locate potential food sources, which include the metabolic breakdown products of plants along with bacteria, fungi and algae (Gilbreath et al., 2013; Gouagna et al., 2012). As an example, baker’s yeast, which is a common component of laboratory rearing diets for mosquito larvae, has been reported to release up to 257 odorant compounds across 13 chemical groups (Alves et al., 2015), many of which have been used in this and other studies of mosquito larvae. Indeed, behavioral assays indicate that yeast odors are extremely attractive to Anopheles larvae (Gonzalez et al., 2019; Xia et al., 2008; Merritt, 1992). Molecular studies of the larval olfactory apparatus, which comprises the antennae along with an apical sensory cone, have characterized 12 putative chemosensory neurons that directly express OR/Orco receptor pairs and Ir76b, one of three IR co-receptors, thereby implicating both OR- and IR-based signaling (Liu et al., 2010; Xia et al., 2008). Moreover, we recently carried out a preliminary RNA-seq-based survey of the larval head, which revealed that multiple IRs—including Ir8a, 25a, and 76b—are significantly expressed (data not shown), further supporting the hypothesis that both ORs and IRs function together in the larval antennae.

In this study, we have completed a detailed cellular investigation of larval olfaction in the Afrotropical malaria vector mosquito *An. coluzzii* by conducting the first comprehensive electrophysiological examination of the mosquito’s larval sensory cone. At first glance, and not surprisingly, when challenged by an odorant panel spanning a wide-range of chemical space, we found distinct responses to a large number of compounds (Figure 1, Supplemental Table S2). It is noteworthy that many of the odorants, such as 1-butanol, 1-octen-3-ol, 1-octanol, 3-octanol, 3-octanone, benzaldehyde, 2-acetylthiazole, and ethylpyrazine, that robustly activate the mosquito’s sensory cone compounds are byproducts of microorganism metabolism, and are likely to function as larval food cues (Alves et al., 2015; Beltran-Garcia et al., 1997; Hung et al., 2015). Furthermore, there is a broad alignment between the robust *in vivo* responses observed here to acetophenone, butylamine, and several ethyl- and methyl-phenols and other compounds that have previously been shown to evoke robust behavioral responses in *An. coluzzii* larvae (Liu et al., 2010; Xia et al., 2008). When these collective responses are visualized as a tuning curve (Figure 1B), it is apparent that the larval sensory cone is a generalist chemosensory appendage that, while biased to alcohols, nevertheless maintains a broad ability to detect a range of chemical classes, including thiazoles, heterocyclics, ketones, aromatics, amines, and carboxylic acids.

The robust complexity of olfactory sensitivity displayed by *An. coluzzii* larvae is further illustrated by an examination of concentration dependency (Figure 2, Supplemental Table S3). While every odorant in the panel elicited dose-dependent responses across five-fold dilutions, albeit with no meaningful activity at the lowest concentration (10^−5^), and several compounds (e.g. 1-octen-3-ol, acetophenone, cyclopentanone, 2-ethylphenol) exhibited saturation kinetics at the highest (10^−1^) odorant dilutions, larval responses to butylamine are notable in terms of intensity and near-exponential kinetics. A survey of odorant tuning curve analyses across several dilutions confirms that the *An. coluzzii* larval sensory cone maintains a generalist sensitivity to environmental cues while displaying a robust sensitivity to several behaviorally active compounds. These include butylamine, which evokes dosedependent effects on *An. coluzzii* larval movements (Liu et al., 2010; Xia et al., 2008); acetophenone, which evokes significant aversive behavior at dilutions as low as 10^−5^; and 4-methylcyclohexanol, which is most attractive to *An. coluzzii* larvae at 10^−4^ dilution (Sih, 1986; Xia et al., 2008). It is noteworthy that these odorants evoked larval behavioral effects that closely mirror their sensory cone response range (Figure 2A/C) as well as their tuning-curve peak prominence across much of that spectrum (Figure 2B). These alignments between larval peripheral neuronal activity and behavior serve to underscore the relationship between these processes.

Despite limitations in precisely discriminating the action potential spikes from individual chemosensory neurons that innervate the larval sensory cone, 12 of which have been previously identified as Orco-expressing ORNs (Xia et al., 2008), their broad amplitude range provided an opportunity to sort them into three distinctive neuronal classes. It is likely that the larval sensory cone spike amplitudes observed here reflect cell body morphology as well as their relative location along the larval antennae, which impacts dendritic length and position (Sih, 1986; Xia et al., 2008). That being said, resolving the collective neuronal responses into discrete bins of A-, B- and C-group sensory neurons further illustrates the dynamic complexity of the larval olfactory system of *Anopheles*. While the breadth, intensity and duration of the B- and C-group responses are more prominent than in A-group neurons (Figure 3), there are nevertheless numerous odorants that selectively activate A-group neurons, occasionally in a sustained manner. These include 3-methylindole (skatole), which is a metabolic byproduct of bacteria (Elgaali et al., 2002; Hubbard et al., 2015; Lindh et al., 2008; Schulz & Dickschat, 2007), fungi (Chen et al., 2014; Tomberlin et al., 2017) and plants (Frey et al., 2000; Ober, 2005; Turlings et al., 1991). It is reasonable to therefore speculate that mosquito larvae might use 3-methyindole as an olfactory cue to locate potential food sources. Indeed, it has been shown to be a potent attractant for Anopheline larvae and specifically recognized by OR2 and OR10, both of which are expressed in the larval antennae (Xia et al., 2008). Several examples of notable B neuron activity are present with, more often than not, a distinctive temporal dynamic consisting of paired 1-s activation intervals followed by a 1-s inhibition. This type of response is characteristic of 1-octen-3-ol and 4-methylcyclohexanol, which have also been characterized as behaviorally active semiochemicals for *An. coluzzii* larvae (Xia et al., 2008). As a whole, the C-group neurons appear more generalist in odor tuning, with activation profiles that are broader and less intense than A- and B-group responses. That said, C-group neurons frequently, as in the case of 1,4-dimethylbenzene and 1,2-dimethoxybenzene stimulation, robustly display immediate inhibitory responses.

A closer examination of the temporal dynamics of neuronal odor coding on the larval sensory cone of *An. coluzzii* brings these interactions into fine detail (Figures 3 and 4). While several of the surveyed odorants narrowly activate either the A- (Figure 4A-D) or B-group neurons (Figure 4E-I), only hexanoic acid evokes a distinct and immediate activation of C-group neurons (Figure 4T). Indeed, this selective activation by C-group neurons is in marked contrast to the largely indifferent responses across the sensory cone to a wide range of other carboxylic acids (Figure 5A), many of which are known to be important in adult host-preference behaviors (Smallegange et al., 2009). Taking this further, we explored the ability of larval sensory cone neurons to discriminate among several other groups of odorants with similar chemical structures, which would be an expected necessary trait for larval navigation in the search for nutrients. Our analyses revealed that, while *An. coluzzii* larval olfactory neurons appeared indifferent to many sulfides and thiols, all three neuron groups differentially contributed to the combinatorial sensitivity to a subset of these compounds (Figure 5B and D). Importantly, the collective involvement of all three groups contributes to the discrimination among aromatic phenols and thiazoles (Figure 5C/D), as well as between the foul-smelling, organosulfur compound thiophenol and the heterocyclic, thiophene, which has a benzene-like odor (Figure 5B).

Attempting to definitively assign a specific OR as well as other chemosensory receptors to the A, B, or C group of larval sensory cone neurons is hindered by the inherently combinatorial relationship between odorants and receptors that underlies the process of odor coding. As previous gene-silencing/targeting studies have implicated that the *Ir76b* co-receptor is involved in Anopheline larva’s response to butylamine (Liu et al., 2010), and IR8a, another IR co-receptor, contributes to the detection of acidic volatiles in *Ae. aegypti* (Raji et al., 2019), it is reasonable to suggest that the responses to butylamine and hexanoic acid presented here are the result of IRNs embedded within A- and C-group larval sensory cone neurons. While the larval ORs have been identified and, in many cases, localized to specific ORNs (Xia et al., 2008) and functionally characterized (Carey et al., 2010; Wang et al., 2010; Xia et al., 2008), it is challenging to be precise. Although these studies identify several larval ORs with relatively narrow tuning breadths, those data also reveal that many odorants seem to promiscuously activate multiple receptors. Nevertheless, a correlative analysis of these responses to tuning OR functionality facilitates a hypothesis that at least a portion of the immediate and robust A-group responses to acetophenone and 2,4,5-trimethylthiazole may be the result *Or6/Orco* and *Or28/Orco* expressing larval ORNs, respectively. However, in light of the paucity of specialized OR-odorant relationships and in the absence of specific tuning *Or* gene-targeting studies, these and other correlations must remain speculative.

The studies presented here show that Anopheline larval olfactory physiology should be viewed as a broadly generalist process that displays intricate dose-dependent and richly discriminating response profiles. Taken together, these data reinforce the premise that Anopheline larvae are complex aquatic insects that benefit from the presence of a versatile olfactory apparatus that provides a robust capacity to navigate and detect a wide range of odorant cues to locate and select nutrients as well as identify predators and other signals they may encounter in their aquatic habitats in order to survive and mature into adults. As such, the larval olfactory system represents a more accessible context in which to study mosquito odor coding and neuronal function, and, perhaps more importantly, as a viable target for the design of novel larval control strategies that could be applied to their relatively restricted aquatic habitats and reduce their potential to develop into adult disease vectors.

---

**Figure S1.**
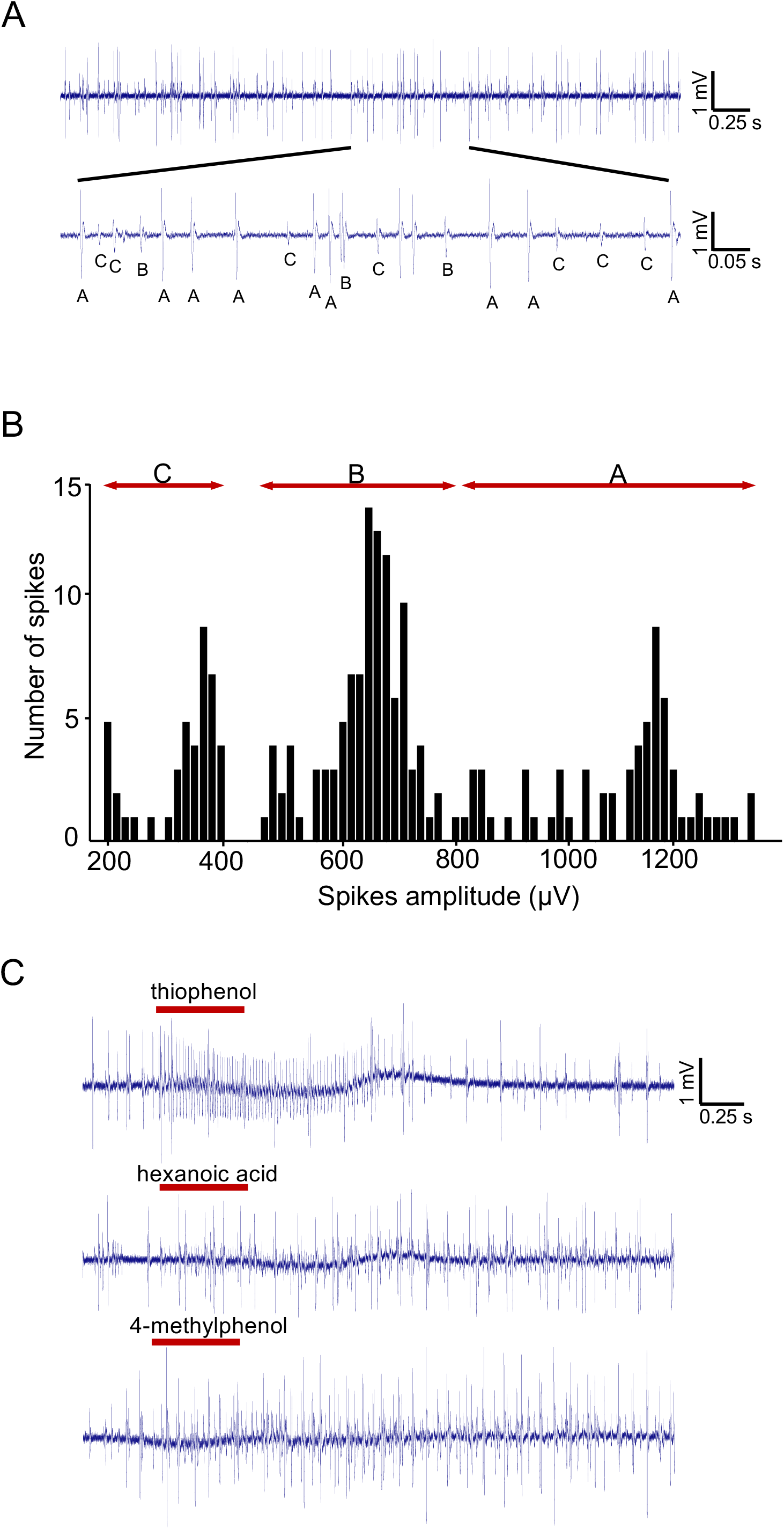
Electrophysiological recording of the *An. coluzzii* larval, antennal sensory cone. (A) Representative signal traces of spontaneous firing in the sensory cone of *An. coluzzii*. Neuronal spikes with different amplitudes are shown. Spikes were classified into three neuron groups (A, B, C), based on amplitude. (B) Multiple clusters using the AutoSpike software for spike sorting are presented. (C) Representative responses of larval chemosensory neurons activated by specific compounds.

**Figure S2.**
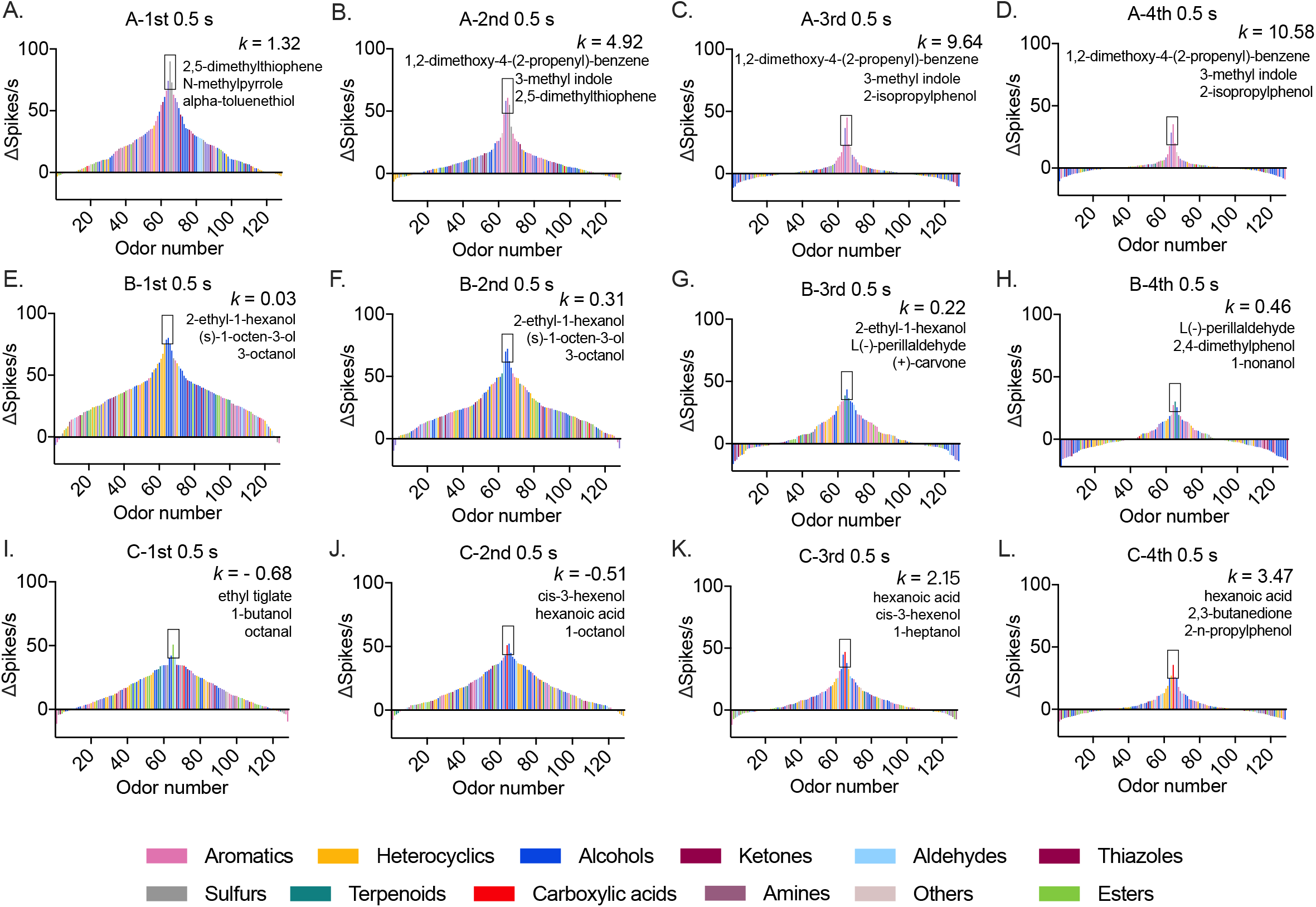
Tuning curves of sorted three neuron groups in response to 128 compounds across a time course of four 0.5-s intervals during and immediately after odor stimulation (n=6-10). The corresponding K values were calculated for each neuron groups in each time interval. The top three odorants contributing to the peak response are listed. All odorants are color coded based on their chemical classes.

**Figure S3.**
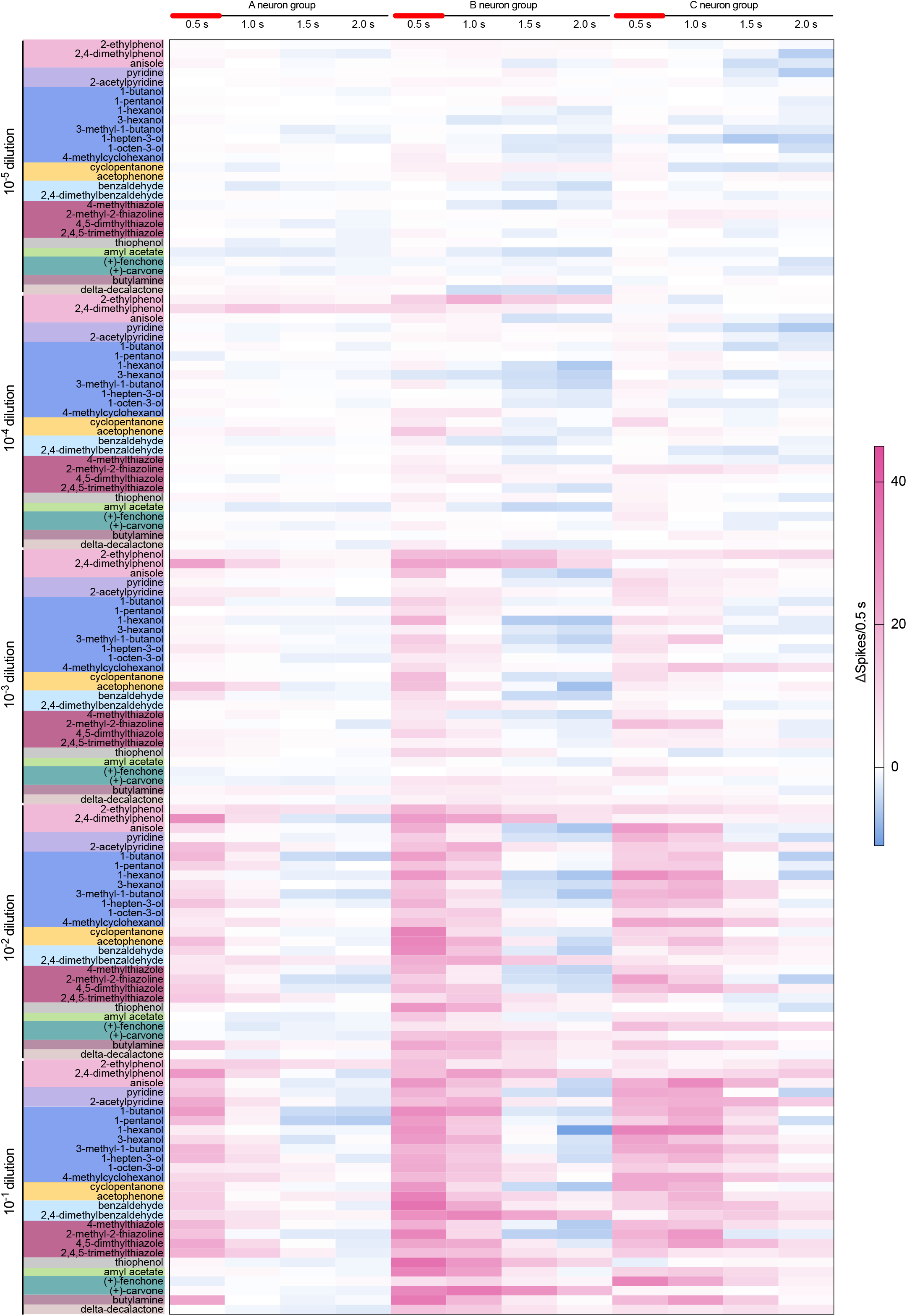
Dose-dependent responses of three neuron groups in response to 27 compounds across five concentrations, ranging from 10^−5^ to 10^−1^ v/v or m/v dilution in paraffin oil or DMSO. n=6-10 for each odorant dilution in each compound.

**Figure S4.**
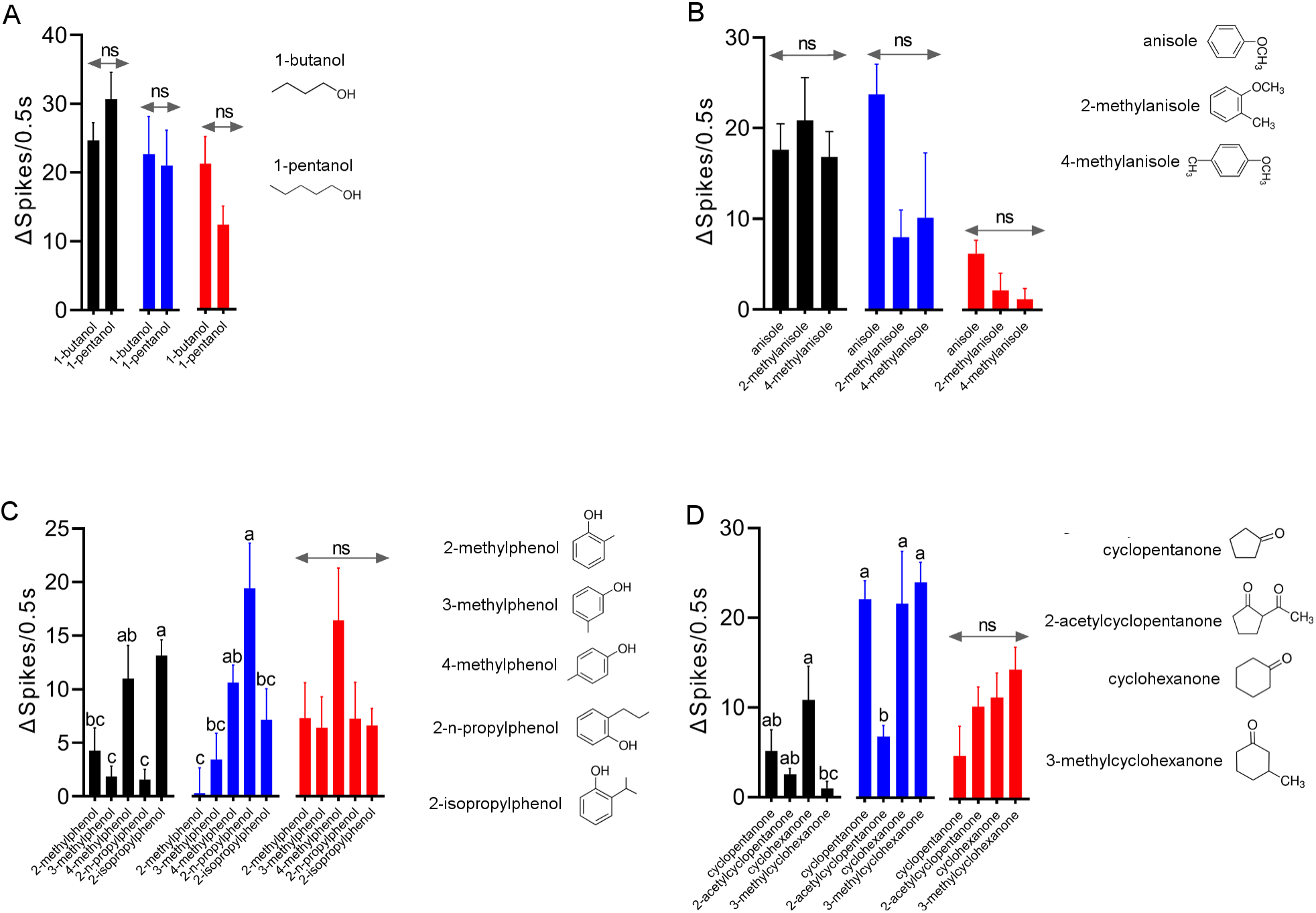
Different responses of the larval, antennal sensory cone to compounds with similar structure (n=6-10). (A) Two alcohol compounds; (B) anisole-derived compounds; (C) isomers of methylphenol; and (D) cyclic ketones. One-way ANOVA Tukey test was applied in the statistical analysis with a p value less than 0.05 indicating significant difference.

Supplemental Table S1. Characteristics of 281 odorants.

Supplemental Table S2. Response profile of the larval, antennal sensory cone to 281 compounds of diverse chemical groups (n=6-10). (a) Mean responses; (b) SEM of responses.

Supplemental Table S3. Dose-dependent responses to a panel of 27 odorants (n=6-10). (a) Mean responses; (b) SEM of responses.

Supplemental Table S4. Response profile of three neuron groups to 128 odors eliciting ≥40 spikes in the sensory cone (n=6-10). (a) Mean responses; (b) SEM of responses.

Supplemental Table S4. Tuning curves of sorted three neuron groups in response to 128 compounds across a time course of four 0.5-s intervals during and immediately after odor stimulation (n=6-10). (a) Mean responses; (b) SEM of responses.

## Contributions

H.S., F.L., and L.J.Z., designed the study. H.S., and F.L. performed experiments. H.S., F.L., and A.B., analyzed data. H.S., F.L., A.B., and L.J.Z interpreted the results, prepared the figures, and wrote the manuscript.

## Acknowledgments

We thank Drs. H. Willi Honegger and Ann Carr, and Mr. Stephen Ferguson for their comments on this manuscript as well as Zi Ye and other members of the Zwiebel lab for critical suggestions and help with data analyses during the course of this work. We also thank Dr. A.M. McAinsh for editorial assistance as well as Zhen Li and Samuel Ochieng for mosquito rearing and technical help. This work was conducted with the support of Vanderbilt University and a grant from the National Institutes of Health (NIAID, AI127693) to LJZ.

